# Somatic copy number variant load in neurons of healthy controls and Alzheimer’s disease patients

**DOI:** 10.1101/2022.05.20.492539

**Authors:** Zeliha Gözde Turan, Vincent Richter, Jana Bochmann, Poorya Parvizi, Etka Yapar, Ulaş Işıldak, Sarah-Kristin Waterholter, Sabrina Leclere-Turbant, Çağdaş Devrim Son, Charles Duyckaerts, İdil Yet, Thomas Arendt, Mehmet Somel, Uwe Ueberham

## Abstract

**Background:** The possible role of somatic copy number variations (CNVs) in Alzheimer’s disease (AD) aetiology has been controversial. Although cytogenetic studies suggested increased CNV loads in AD brains, a recent single-cell whole-genome sequencing (scWGS) experiment, studying frontal cortex brain samples, found no such evidence. Here we readdressed this issue using lowcoverage scWGS on pyramidal neurons dissected using laser capture microdissection (LCM) across five brain regions: entorhinal cortex, temporal cortex, hippocampal CA1, hippocampal CA3, and the cerebellum.

**Results:** Among reliably detected somatic CNVs identified in 1301 cells obtained from the brains of 13 AD patients and 7 healthy controls, deletions were more frequent compared to duplications. Interestingly, we observed slightly higher frequencies of CNV events in cells from AD compared to similar numbers of cells from controls (4.1% vs. 1.4%, or 0.9% vs. 0.7%, using different filtering approaches), although the differences were not statistically significant. We also observed that LCM-isolated cells show higher within-cell read depth variation compared to cells isolated with fluorescence activated cell sorting (FACS), which we argue may have both biological and technical causes. Furthermore, we found that LCM-isolated neurons in AD harbour slightly more read depth variability than neurons of controls, which might be related to the reported hyperploid profiles of some AD-affected neurons. We also propose a principal component analysis-based denoising approach that significantly reduces within-cell read depth variation in scWGS data.

**Conclusions:** We find slightly higher somatic CNV frequencies in the brains of AD patients, and higher sequencing coverage variability, although the effects measured do not reach statistical significance. The results call for improved experimental protocols to determine the possible role of CNVs in AD pathogenesis.

## Introduction

**<u>A</u>**lzheimer’s **<u>d</u>**isease (AD) is a neurodegenerative disease of multifactorial aetiology, with numerous genetic and environmental factors each explaining a small proportion of variance in disease onset and progression (1). One of the less-studied potential contributors is somatic **<u>c</u>**opy-**<u>n</u>**umber **<u>v</u>**ariations (CNVs) in neurons, which can include the gain or loss of whole chromosomes (aneuploidy) or of chromosomal segments. It is generally accepted that mature neurons in healthy brains can carry somatic CNVs, but their frequency is uncertain. Early studies estimated aneuploid neuron frequencies between 4% to 40% in neurotypical brains (2–4), while analyses using **<u>s</u>**ingle-**<u>c</u>**ell **<u>w</u>**hole-**<u>g</u>**enome **<u>s</u>**equencing (scWGS) estimated aneuploid neuron frequencies at <1% (5). Beyond aneuploidy, recent scWGS studies also estimated CNV-carrying neurons at around 30% in young adults and 10% in old adults (6).

Over the last two decades, a number of **<u>f</u>**luorescence **<u>i</u>**n **<u>s</u>**itu **<u>h</u>**ybridization (FISH) and cytogenetic-based studies investigated CNV frequencies in AD and healthy control brains (2, 7–12). Several of these reported extra copies of chromosomes in the AD brain (7–12). This, in turn, implies that the chromosomal imbalance might contribute to AD pathogenesis via altered gene expression levels. An example of such imbalance is seen in individuals with Down’s syndrome (DS); carrying an extra copy of chromosome 21 appears to facilitate aggregation of amyloid-*β* (A*β*) plaques in the brains of DS individuals similar to the AD phenotype (9, 13, 14).

There are various explanations for why post-mitotic neurons in AD brains could carry high frequencies of somatic CNV (15). According to one view, the high CNV burden in the AD brain originates from neurogenesis in the embryonic period. This excessive somatic mutation may be pathogenic and manifest itself as increased AD risk during ageing (16). However, Abascal *et al*. recently showed that somatic mutation (single nucleotide change or indel) accumulation in cells with mitotic capacity and in post-mitotic neurons follow similar trajectories. That is, mutational processes (possibly also including CNVs) appear to occur in a time-dependent manner rather than being division-dependent (17). Accordingly, CNVs in AD brains may have accumulated during their lifetime. However, this scenario also appears inconsistent with the observation that CNV-bearing neuron frequencies decrease from young to old adulthood (6). Another view suggests that AD itself might cause dysregulation in neurons, and AD-affected mature neurons might re-enter the cell cycle, resulting in increased CNV load (8, 18), which may then be eliminated at later stages of AD, thus causing neurodegeneration (10).

Over the last decade, advances in **<u>n</u>**ext-**<u>g</u>**eneration **<u>s</u>**equencing (NGS) technologies gave fresh impetus to somatic CNV analyses by allowing variants to be determined at the single-cell level (19). In one such study, van den Bos and colleagues used scWGS to compare the prevalence of aneuploidy in neurons from healthy control and AD patients (5). Analyzing 1482 neurons from 10 AD patients and 6 control individuals, the authors reported aneuploid prevalence at 0.7% and 0.6% for control and AD neurons, respectively, and concluded that aneuploid cells are not more common in the AD brain.

These findings by van den Bos and colleagues implied that CNVs may have no relationship to AD pathogenesis, in contrast with earlier finds from FISH and cytometry. However, the study by van den Bos and colleagues had a number of limitations. One was that the authors only estimated aneuploidy (full chromosome gain or loss), while large CNVs, which could also contribute to pathogenesis, remained uncharacterized. Another limitation was that only one brain region was examined, the frontal cortex, while atrophy of the medial temporal lobe and specifically the hippocampus is generally considered to be a strong predictor of AD (20). The study did not distinguish among neuron types that may carry differentially sensitivity to AD (21). Thirdly, the study discarded a large fraction of cells (39%) for showing high within-cell variability in genome coverage, although it was unclear to what extent these represented pure technical error versus cells with complex karyotypes. Fourthly, only NeuN positive neurons were included, which substantially restricts the significance of this study due to different reasons: *(1)* Recently, up to 30% of cortical neurons have been reported being NeuN-negative following diffuse brain injury and which are more vulnerable to membrane disruption (22), a process recently associated with AD (23, 24). *(2)* Considerable or even complete loss of NeuN immunoreactivity was also reported for neurons affected by ischemic insults (middle cerebral artery occlusion) without significant cell loss (25) or in neurons that just entered the cell death process (26). Interestingly, these neuronal populations are of special interest because energy and nutritional deficiency and cell loss are essential characteristics of the AD brain (27). *(3)* The intensity of NeuN staining is reported to be lower in AD samples (28), and further *(4)* due to many NeuN negative cortical neurons in FTLD-TDP (frontotemporal lobar degeneration with TDP-43 inclusions) patients, Yousef *et al*. suggested NeuN staining as an indicator of healthy neurons (29). However, if NeuN reflects a neuron’s health, any selection of NeuN positive cells would lead to a substantial bias for studying any neurodegenerative disease.

These methodological issues could potentially explain the discrepancies between the findings by van den Bos *et al*. and those based on FISH and cytogenetic studies (7–12). Notably, a recent technical comparison between FISH and scWGS using mock aneuploid cells reported a tendency of the latter to severely underestimate aneuploidy (30). It is thus possible that both neurons with CNV and nuclei thereof display altered physico-chemical properties. This may result in selection bias against abnormal nuclei with high CNV loads when using the **<u>f</u>**luorescence **<u>a</u>**ctivated **<u>c</u>**ell/**<u>n</u>**uclei **<u>s</u>**orting (FACS, FANS) isolation method (exerting mechanical stress (31)) and high hydrodynamic pressure (32), applied by van den Bos and colleagues, and artificially inflate euploidy frequencies. Moreover, besides restriction to NeuN positive cells, usage of only intact nuclei could preclude or bias AD neurons with nuclear envelope stress or rupture (33).

These observations call for additional data and approaches to tackle this issue. Accordingly, here we generated and analyzed scWGS data to establish the frequency of CNVs (both full chromosome aneuploidies and sub-chromosomal CNVs) in five different brain regions that differ in vulnerability to AD (34). We employed two different single-cell isolation methods, **<u>l</u>**aser **<u>c</u>**apture **<u>m</u>**icrodissection (LCM) and FACS, to isolate neuronal nuclei. LCM, despite being technically challenging, has the advantages of allowing for specific neuron types to be chosen, and being neutral towards normal and abnormal nuclei. We further employed a principal component analysis-based denoising approach to eliminate false positive CNV calls that might result from either systematic experimental biases or repetitive regions in the human genome. Finally, we analyzed published datasets to replicate our main results and check the sensitivity and specificity of our bioinformatics pipeline.

## Results

### Summary of the dataset

We used scWGS to determine the frequency of CNVs in the temporal cortex, hippocampal subfields **<u>c</u>**ornu **<u>a</u>**mmonis (CA) 1, hippocampal subfields cornu ammonis (CA) 3, **<u>c</u>**ere**<u>b</u>**ellum (CB) and **<u>e</u>**ntorhinal **<u>c</u>**ortex (EC) of 13 AD patients and 7 age-matched healthy controls (Fig. 1A-C, Additional file 1: Table S1). The Braak stages of AD patients ranged between III and VI (Fig. 1D). Neuronal nuclei were isolated using either FACS (sorted with propidium iodide, *n* = 12) or LCM (sorted with cresyl violet, *n* = 1552), the latter performed on frozen brain slices (see Methods). LCM-isolated non-neuronal “blank” regions were used as negative control (*n* = 10). The LCM method, although more difficult to implement than FACS, was chosen to ensure the selection of nuclei of pyramidal neurons for sequencing, known to be particularly sensitive to AD (21). For technical comparison, neurons of a single individual were collected both using FACS (*n* = 12) and LCM (*n* = 64) (see Methods). scWGS libraries were prepared using GenomePlex whole-genome amplification and specific adapters were inserted using Phusion^®^ PCR. Paired-end reads were mapped to the human reference genome, followed by stringent filtering to obtain uniquely mapped reads. This resulted in a median of 276,446 reads, corresponding to a coverage of 0.006X per LCM-isolated cell (range: [133 - 1,909,016] reads and [0.000003X - 0.04X] coverage) (Fig. 2A-B).

**Fig. 1.**
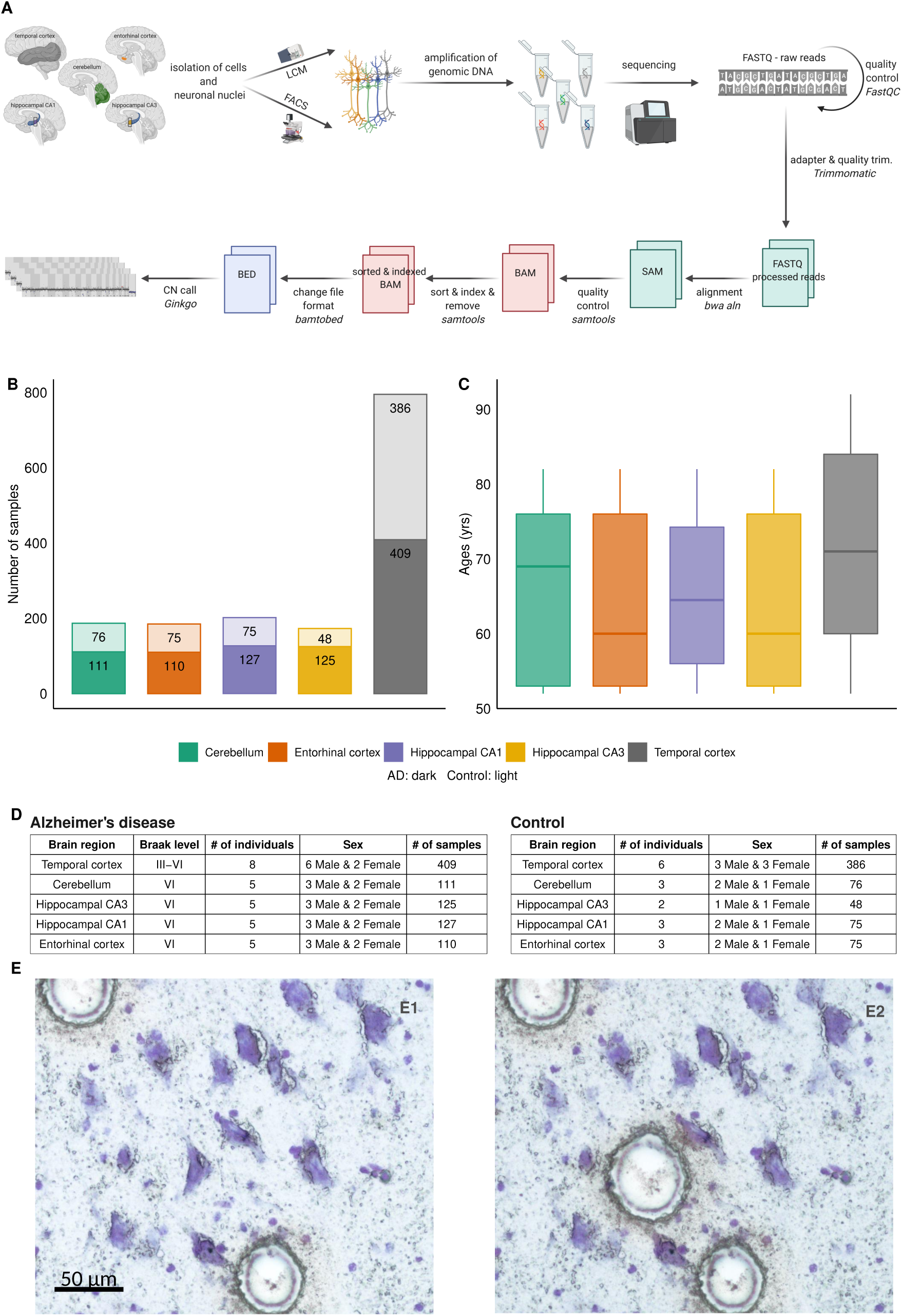
Schematic of the workflow and information about the samples. **A** The pipeline of NGS data analysis and CNV detection. **B** Bar plot showing the number of cells that have been sequenced for each brain region. **C** Boxplot showing the age distribution of samples for each brain region. **D** The tables contain additional information about the sex, diagnoses and Braak level of the samples. **E** Images from a frozen hippocampal brain slice stained with cresyl-violet showing a pyramidal cell before *(E1)* and after *(E2)* laser capture microdissection-based isolation process using the PALM device. Circles in E1 indicate positions where two pyramidal cells have already been isolated just prior to the picture being taken. Bar, 50 µm.

**Fig. 2.**
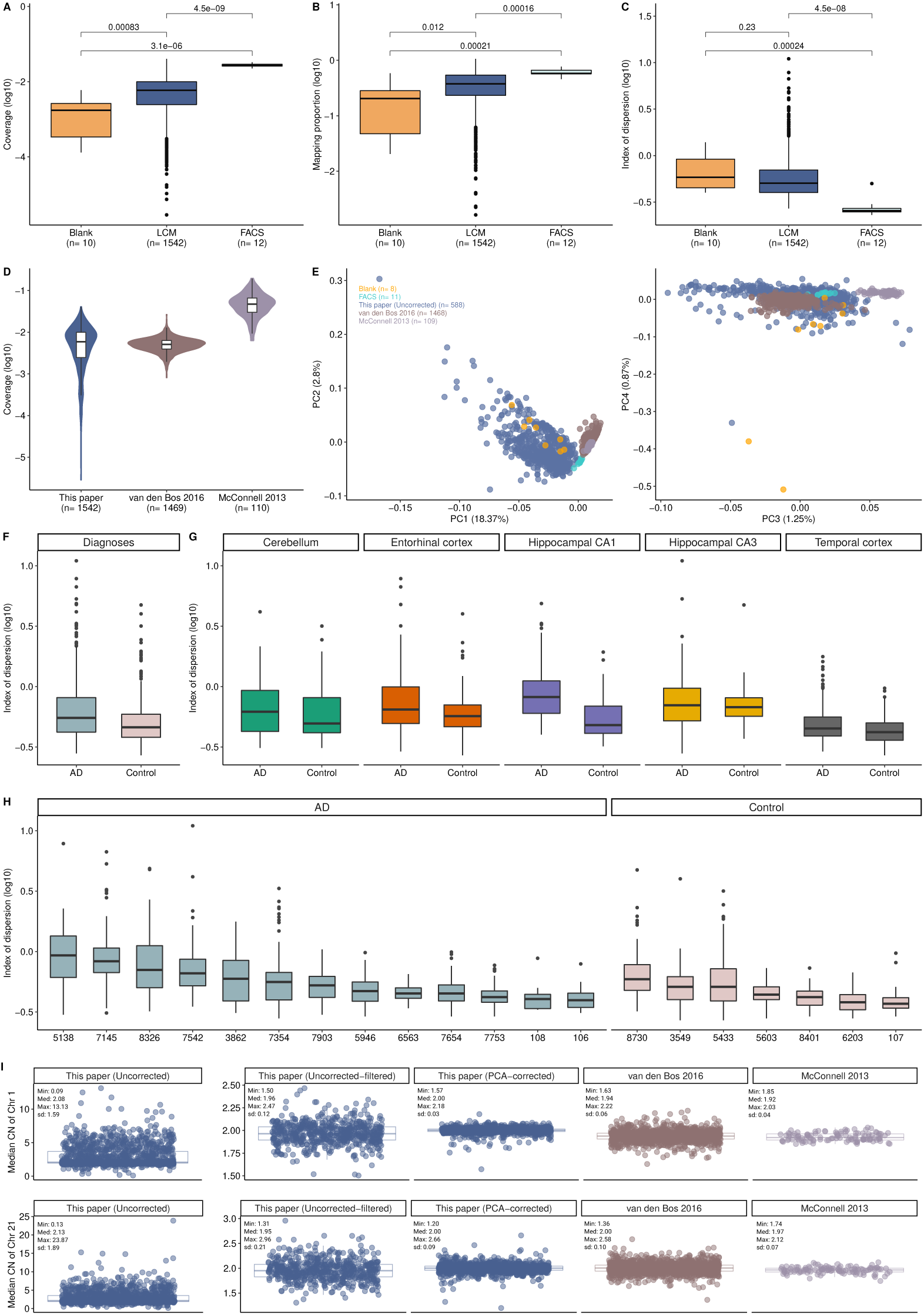
Comparison between different cell isolation methods and published datasets. Box plots showing the distribution of coverage **A**, mapping proportion **B**, index of dispersion **C** among FACS-isolated, LCM-isolated and LCM-isolated blank samples. *P* -values were calculated using Kruskal–Wallis test among groups and Wilcoxon rank-sum test between groups. **D** Violin plot showing the distribution of coverage among different datasets. This study, including only LCM: blue; van den Bos 2016: brown; McConnell 2013: purple. **E** A principal components analysis (PCA) was performed using the normalized read counts across autosomal bins (*n* = 5243) in published datasets and this study. Because they dominated the PCs, cells deviating from the [1.9-2] range were not included in the analyses. The number of cells for each dataset were indicated on the plot. X-axes illustrate PC1 and PC3 that explain 18.4% and 1.3% of the total variance, respectively. Y-axes show PC2 and PC4 that explain 2.8% and 0.9% of the total variance, respectively. The distribution of index of dispersion (“IOD”) for LCM-isolated cells (*n* = 1542) according to **F** diagnoses (Alzheimer’s disease (“AD”), control), **G** brain regions (hippocampal subfields CA1 (“Hippocampal CA1”), hippocampal subfields CA3 (“Hippocampal CA3”), Entorhinal cortex, Cerebellum, Temporal cortex). For each brain region, we tested whether AD diagnosis was predictive of IOD using a linear mixed-effects (lme) model (see Methods). Individuals were added as a random factor. Across all tested brain regions, differences were only marginally significant (*p* = 0.069). **H** The distribution of IOD across individuals (*n* = 20). Box plots were ordered by the median. Y-axes illustrate the IOD values on the log10 scale. **I** Boxplots showing the distribution of median CN of chromosome 1 (chr1, upper part of the figure) and chromosome 21 (chr21, lower part of the figure) across bins (*n* = 440 and *n* = 68 for chr1 and chr21, respectively). Each point corresponds to the median CN of each cell. Minimum (“Min”), median (“Med”), maximum (“Max”) and standard deviation (“sd”) of each distribution were shown on the boxplot. Cells that deviated from the [1.9-2] range were excluded from the analyses to be consistent with our filtering criteria (except for the uncorrected datasets). This study [Uncorrected (*n* = 1337), Uncorrected-filtered (*n* = 588), PCA-corrected (*n* = 1301)]: blue; van den Bos 2016 (*n* = 1468): brown; McConnell 2013 (*n* = 109): purple.

CNVs were predicted using the *Ginkgo* algorithm, which uses circular binary segmentation (CBS) to estimate deletion or duplication events (35). Negative controls (*n* = 10) and FACS-isolated neurons (*n* = 12) were analyzed separately and are not included in the main results. *Ginkgo* was run on our dataset with *n* = 1542 cells, while in parallel, two published scWGS datasets were also analyzed: one by van den Bos and colleagues (“van den Bos 2016”), comprising *n* = 1469 cells from healthy and AD brains (median coverage 0.005X), and another by McConnell and colleagues (“McConnell 2013”), comprising *n* = 110 cells from healthy brains (median coverage 0.047X) (Fig. 2D) (5, 36). Note that the van den Bos 2016 dataset includes only 61% of cells produced in that study, because data from cells filtered for high noise levels were not published and thus could not be included here.

### LCM-isolated cells show a high frequency of depth variability

We first evaluated the sensitivity and specificity of our bioinformatics pipeline on scWGS data using trisomy-21 in DS and monosomy-X in males in published data. Analyzing *n* = 34 neuronal nuclei from DS individuals (5), trisomy-21 was correctly predicted across all samples without any false positive or false negative calls. In addition, monosomy-X was accurately predicted in 94.2% (338 of 359) of cells from males across the two published datasets (5, 36).

*Ginkgo* includes an algorithm that uses the distribution of read depth across the genome to infer the average DNA copy number of each cell, which is estimated within a range of 1.5 to 6. It would be expected that the majority of human neurons would carry on average two copies of each autosome. Indeed, applying *Ginkgo* on the two published datasets, we found that for 99.9% (1577 of 1579) of cells the estimated average copy number lies within [1.9-2]. Using the same algorithm on our dataset, however, only 45% (687 of 1542) of the cells had average copy numbers estimated within the [1.9-2] range; i.e. 55% were non-euploid. Although hyperploid neurons have been described in control brains at ∼ 10% frequency using FISH (10), the observed non-euploidy estimates suggest that our dataset carries particularly high levels of variability in read depth. These differences, in turn, could be related to the LCM protocol used, as the published scWGS experiments had used FACS.

To investigate this possibility, we compared the quality metrics of cells we had collected using FACS or LCM for this study. These metrics were mapping proportion (the number of mapped reads/ the total number of reads), coverage, and index of dispersion (IOD, the ratio between the variance of read coverage and the mean). FACS-isolated cells had higher sequencing coverage and mapping proportions than the LCM-isolated ones (Wilcoxon two-sided rank-sum test, *p <* 0.0001 and *p <* 0.001 for coverage and mapping proportion, respectively) (Fig. 2A-C). In addition, FACS-isolated cells had low IOD values, indicating less variation in sequence depth than the rest of the samples (Kruskal–Wallis test, *p* = 1.5*e −* 07) (Fig. 2C). We note that the higher noise observed in LCM data was not solely due to higher genome coverage, as the FACS-based data from the van den Bos 2016 dataset had a median coverage comparable to ours (0.005X vs. 0.006X), but did not show comparable variability as in our LCM data (Fig 2D). These differences in IOD between LCM and FACS could be potentially explained by the higher sensitivity of the LCM procedure to experimental noise, compared to FACS. Alternatively, they could partly represent abnormal nuclei selected out in FACS but captured by LCM.

We next investigated the possibility that underlying variation may be caused by technical and/or biological factors. For this, we used a generalized linear mixed model (GLMM) to explain IOD (the response variable) per LCM-isolated cell (*n* = 1542) as a function of diagnosis (AD vs. control), genome coverage, and brain region as fixed factors, and individual as a random factor (see Methods; Fig. 2F-H). We found that coverage has a significant negative effect on IOD (*z* = − 21.06, *p <* 0.0001). Compared to the cerebellum, the region least affected by neurodegenerative diseases (Xu et al. 2019), we found a significantly high IOD for the entorhinal cortex (*z* = 2.61, *p <* 0.05), hippocampal CA1 (*z* = 3.34, *p <* 0.001) and hippocampal CA3 (*z* = 3.75, *p <* 0.001), but not for the temporal cortex (*z* = − 0.28, *p* = 0.78). Finally, neurons from control individuals have slightly less IOD than AD patients (*z* = − 1.93, *p* = 0.054). This result might suggest a tendency for neurons of AD patients to carry more variable DNA content and is consistent with cytometry analyses reporting a high occurrence of hyperploid neurons in the AD brain (10). Although these findings imply a role of biological factors in read count variation within cells, it still remains possible that confounding technical factors influence our data. Given this uncertainty about the source of variability, we continued the analyses by filtering our dataset to remove the most variable cells.

### No significant difference in CNV frequency between AD and control in the “uncorrected-filtered” dataset

We then used *Ginkgo* to call CNV events from “uncorrected-filtered” dataset (*n* = 882 cells from 13 AD patients, and *n* = 660 cells from 7 healthy controls). We found 19,608 events in 882 cells from AD patients (22.2 per cell), and 14,844 events in 660 cells from healthy controls (22.5 per cell). We then tested the observed frequency difference between AD and control using a GLMM with a negative binomial error distribution (see Methods). The response variable (the frequency of CNVs) was predicted using a combination of fixed factors, including diagnoses, chromosomes, brain regions, sex and coverage (Fig. 4D). The individual effect was added as a random factor. We found no statistically significant difference between AD and control across all tested combinations (GLMM, *p ≥* 0.17; Additional file 3: Table S3).

CNV estimation from low coverage scWGS data is known to be highly sensitive to technical noise, and a large proportion of the called CNV events likely represent false positives. We thus decided to filter both cells and CNV events in our dataset to obtain a more reliable dataset (6, 37, 38). We started by removing the most highly variable cells among the LCM-isolated ones (*n* = 1542) using the following criteria. First, 13% (205 of 1542) of the cells with a low number of reads (*<* 50, 000) were discarded from the analysis (see Methods). Second, as most cells are expected to be diploid, and also given that the *Ginkgo*-estimated **<u>c</u>**opy **<u>n</u>**umber (CN) profiles of 99% of cells in the McConnell 2013 and van den Bos 2016 datasets were observed to lie between [1.9-2], we excluded those cells with CN values beyond this range (54% excluded, 726 of 1337). Third, we filtered out 23 of the remaining 611 cells (4%) that showed extreme CNV intensity, which we defined as three or more chromosomes of a cell carrying predicted CNVs that cover *>*70% of their length (Fig. 4A). Information about the remaining cells (*n* = 588) is provided in Additional file 2: Table S2 and Additional file 4: Fig. S1.

From these 588 cells, we called 3521 CNVs (∼ 5.9 events per cell) in the uncorrected data, which we call the “uncorrected-filtered” dataset. We further applied a number of conservative filtering criteria to remove potential false positives: *(1)* We only included megabase scale CNVs (≥ 10 Mb), considering that detection of small events with low coverage data will be unreliable. *(2)* We limited the analyses to 1-somy and 3-somy events, assuming that most somatic CNVs involving chromosomes or chromosome segments would involve loss or duplication of a single copy. *(3)* We only included CNVs with unique boundaries across all analysed cells, assuming that somatic CNV breakpoint boundaries should be generally randomly distributed across the human genome. *(4)* We removed CNVs on the proximal portion of the chr19 p-arm, where frequently observed duplications were previously reported as low coverage sequencing artifacts (39). *(5)* To ensure the reliability of the CNV signal, we calculated a standard *Z*-score for each CNV that reflects the deviation in read count distribution in that region compared to the rest of the cell (called *Z*_1_, see Methods), and only accepted CNVs with absolute values of *Z*_1_-scores ≥ 2. *(6)* We reasoned that read counts in a real CNV should be closely clustered around expected integer values (*e*.*g*. 1 or 3). To assess this, we calculated a *Z*-score for the deviation from the expectation (called *Z*_2_), and only accepted events with absolute values of *Z*_2_- scores ≤ 0.5 (see Methods, Fig. 4A, Additional file 4: Fig. S3).

After CNV filtering, we found 12 CNV events across 295 cells in 13 AD individuals and 4 CNV events across 293 cells in 7 controls. Among the 295 pyramidal neurons analyzed from the 13 AD patients, we found 10 deletions (3.39% per cell) and 2 duplications (0.68% per cell). These events ranged in size from about 10.14 to 77.01 Mb (median: 19.31 Mb) and were observed in the temporal cortex and the entorhinal cortex. Of the 293 neurons from 7 control brains, 1 deletion (0.34% per cell) and 3 duplications (1.02% per cell) were detected in the temporal cortex with a size range of 10.81 to 54.67 Mb (median: 14.51 Mb). Again testing the CNV frequency differences between AD and control brains using a GLMM, we found no statistically significant effect (GLMM, p ≥ 0.88) (Fig. 4B, Additional file 2: Table S2, Additional file 3: Table S3).

### A PCA-based denoising approach minimizes within-cell depth variability

To gain further insight into within-cell variability in our dataset (the uncorrected-filtered version) compared to the two published scWGS datasets, we calculated the median CN of chr1 and chr21 (the largest and smallest chromosomes) across all three datasets. We still found conspicuously higher within-cell variation in our dataset, despite having discarded highly variable cells (Fig. 2I). We then used the autosomal normalized read counts to perform a PCA on the uncorrected-filtered data and published datasets. We also included blank (negative control) samples and FACS-isolated cells to illustrate how reads counts from these two groups relate to others. According to the PCA, LCM-isolated uncorrected-filtered data and blank samples were separated from the published datasets and FACS-isolated cells (Fig. 2E). This result might also highlight distinct profiles of LCM-isolated cells.

We then sought an approach that could reduce this elevated within-cell variability in read depth, assuming it is of technical origin and possibly related to the LCM procedure. Experimental biases could involve cross-contamination across cells during isolation, or biases that arise during DNA amplification. Although the former should be mainly random, the latter may follow systematic patterns, such as some chromosome segments being more or less prone to be amplified.

We thus devised a procedure for removing putative patterns of systematic read depth variation across cells (see Methods). The algorithm starts by choosing a focal cell x in the dataset, and calculating principal components (PCs) from the normalized read counts per autosome across the rest of the cells (except cell x). It then collects all PCs explaining ≥ 90% of the variance. Treating these as representatives of systematic variation, it removes their values from the normalized read counts of cell x using multiple regression analysis. These steps are performed on all cells individually, creating a denoised dataset. The final dataset contains residuals from the multiple regressions instead of the normalized read counts. Notably, this procedure should remove experimentally-induced variation in read depth shared among cells, and also any recurrently occurring somatic CNVs. Rare somatic CNVs, instead, would be mostly unique to each cell and randomly distributed in the genome, and thus would not be affected.

After filtering cells with a low number of reads (*n* = 205) and denoising our dataset with this approach, CN and CNV prediction were performed using Ginkgo. We further compared the results between the PCA-corrected and “uncorrected-filtered” datasets. Examples of cells having “noisy” profiles before and after correction are shown in Fig. 3A-C, which suggests a dramatic reduction in within-cell variability. Beyond visual inspection, we also analyzed three statistics. First, we studied the CN profile of cells after PCA correction. We found 97% (1302 of 1337) now lie between 1.9 and 2 (Fig. 4C). This result is comparable to the two published datasets described above and much higher than uncorrected data (45%). Second, we calculated the number of CNV events per cell (sum of the number of CNV/ number of cells) across datasets. In the van den Bos 2016 and McConnell 2013 datasets, we estimated 5.6 and 8.1 CNVs per cell, respectively (Fig. 3E). In our dataset, in the uncorrected version, we found 23.9 CNVs per cell, in the “uncorrected-filtered” data 6.0 CNVs per cell, and in the PCA-corrected data, we estimated on average 1.0 CNV event per cell. The correction leads to lower CNV estimates in our data, which is more conservative and possibly more realistic than the higher estimates without correction. Third, we estimated the standard deviation in CN among cells for chr1 and chr21. For chr1 and chr21, the standard deviations in the PCA-based data were 4 and 2.3 times lower than in the “uncorrected-filtered” data, respectively, and comparable to CN standard deviations in the two published datasets.

**Fig. 3.**
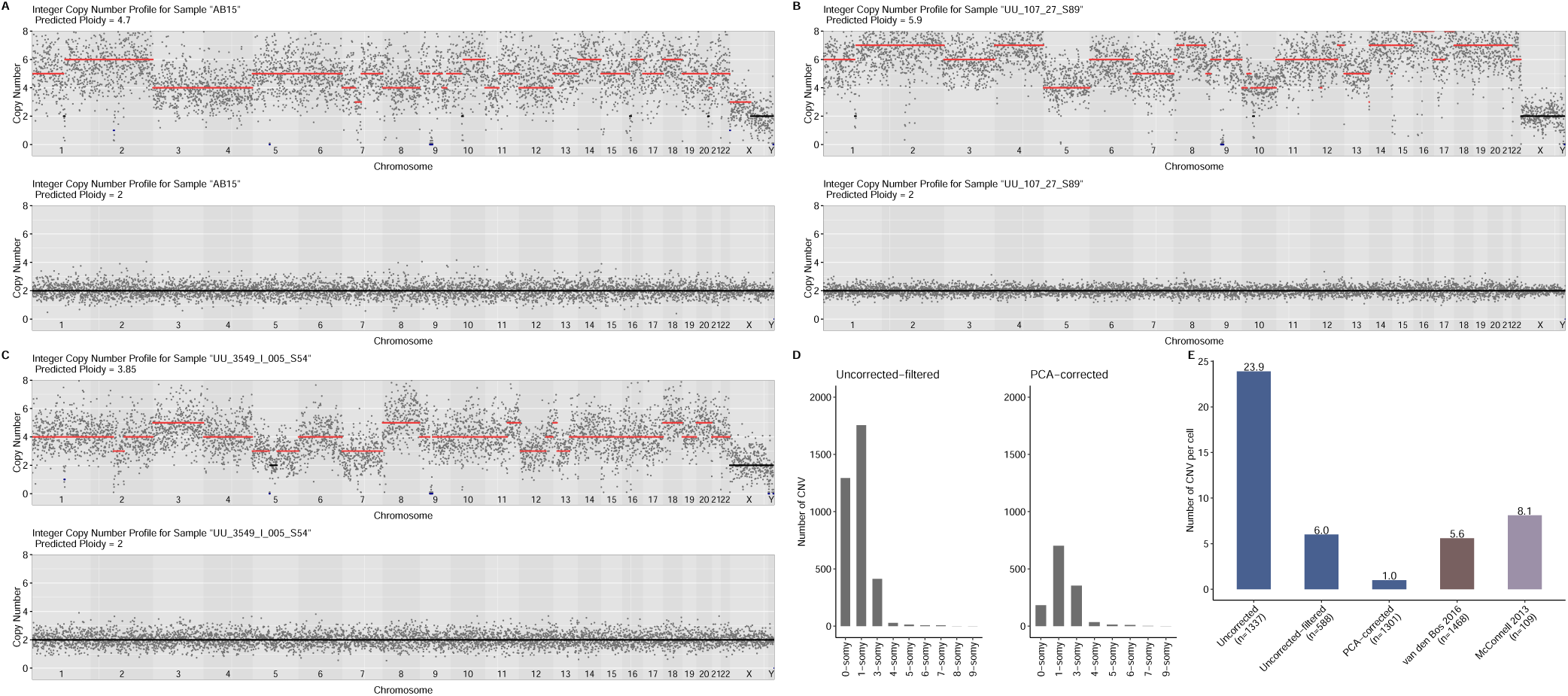
A PCA-based denoising approach helps eliminate technical noise. **A-C** Examples of CN estimates of cells using uncorrected data (upper panels) and using data after PCA-based correction (lower panels). The x-axes show chromosomes and the y-axes show the CN profile of chromosomes estimated by *Ginkgo*. Each grey dot represents the scaled and normalized read counts per bin. Amplifications (CN*>*2) are shown in red; deletions (CN*<*2) in blue; disomy (CN=2) in black. Our denoising approach did not work well for three cells and the CN profiles of them are shown in Additional file 4: Fig. S2. **D** Distributions of the number of CN before (*n* = 3, 521) and after correction (*n* = 1298) across autosomes. For illustration purposes, the bar plot includes up to 10-somy. **E** Bar plot showing the number of CNVs per cell across datasets. The datasets from our study include cells from both AD and control (Uncorrected, Uncorrected-filtered, PCA-corrected): blue; van den Bos 2016 (including cells from both AD and control): brown; McConnell 2013: purple). The number of cells that were used to calculate CNVs per cell was shown on the X-axis label.

### Subchromosomal CNVs are enriched in deletions in the PCA-corrected data

Based on these three statistics, we decided to study this PCA-corrected version of our datasets. For downstream analysis, we further eliminated cells that deviated from the ploidy range of [1.9-2] (2.6%, 35 of 1337) or showed extreme CNV intensity (0.08%, 1 of 1302) (Fig. 4A). We thus created a denoised dataset of 1301 pyramidal neurons from 20 individuals.

Estimating CNVs in this dataset using Ginkgo, we found 1298 CNVs in total (∼ 1 event per cell). To remove false positives, we also performed the same CNV prediction and downstream analyses on our PCA-corrected data (Additional file 4: Fig. S3). After these steps, we found a total of 9 deletion events (0.7% per cell) and 1 duplication event (0.08% per cell) across 1301 cells in 20 individuals among all tested brain regions (except for the hippocampal CA1 where no CNV event was found). This excess of deletions is unexpected under the null hypothesis of equal expectation of duplication and deletions (two-sided binomial test *p* = 0.021), but consistent with previous observations of more deletions than duplications among somatic mutations (6, 36, 38, 40).

### No significant difference between AD and control after PCA-correction or in the van den Bos dataset

Studying CNV frequencies with respect to diagnosis, we found 6 CNV events across 688 cells in 13 AD individuals and 4 CNV events across 613 cells in 7 controls (Fig. 4E). Performing the formal test for the hypothesis of AD versus control differences with this data, we again found no significant difference between the groups (GLMM, p *≥* 0.80; Additional file 3: Table S3). Information about the CNVs and cells can be found in Additional file 2: Table S2.

**Fig. 4.**
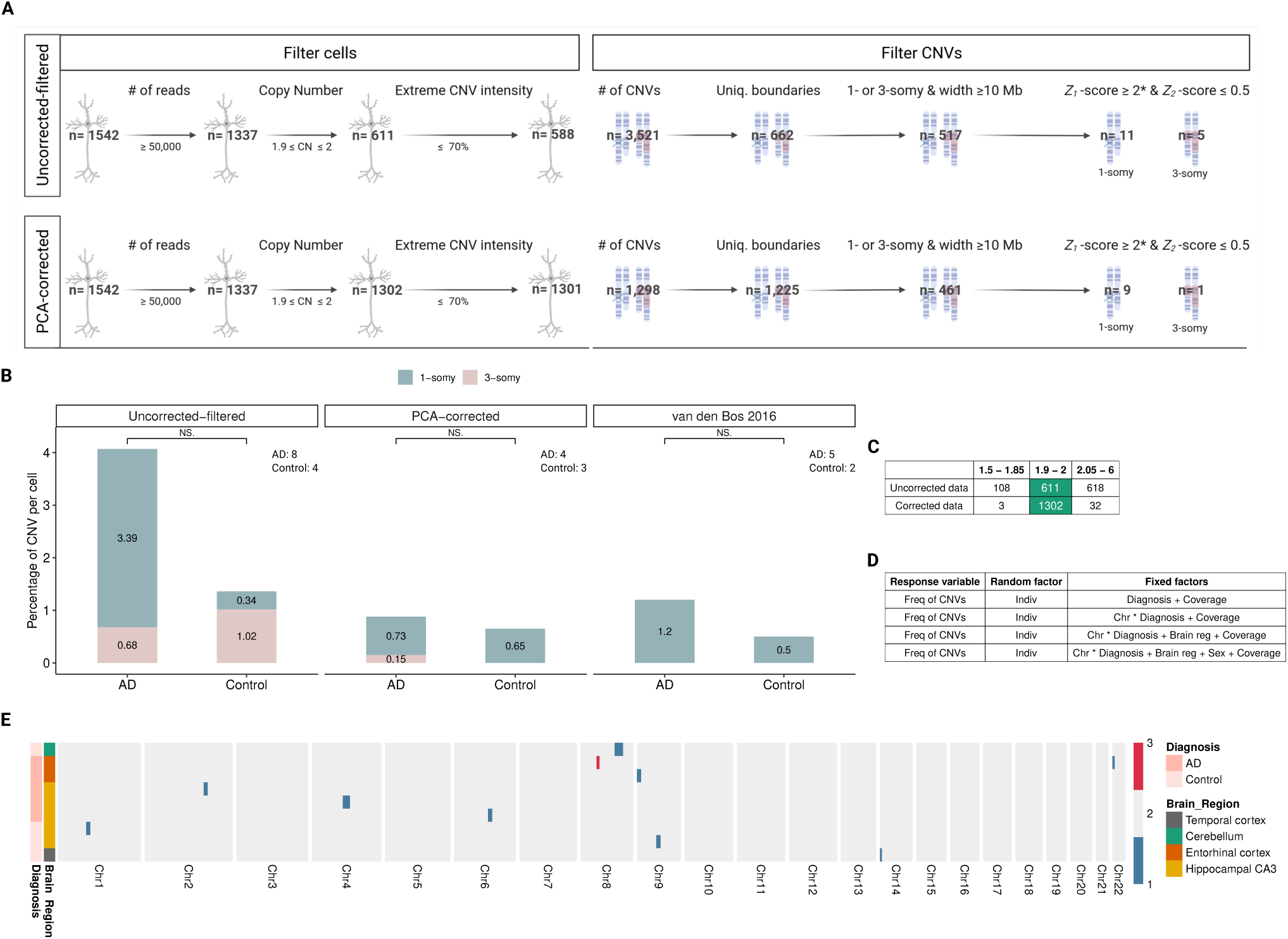
Genome-wide distribution of predicted CNVs in neurons. **A** Overview of cell and CNVs elimination steps in the uncorrected and corrected data. *We discarded CNVs if their breakpoints were within the three base pairs window around each other. **B** Bar plots represent the number of CNVs per cell for AD and control groups. Blue area: 1-somy (deletions); pink area: 3-somy (duplications). The number of unique individuals with at least one cell identified with a reliable CNV is indicated within the panel. **C** The table shows the ploidy levels of cells in corrected and uncorrected versions of the data. **D** The table shows several combinations of fixed factors that were used to test the difference between AD and control in the frequency of CNVs (“Freq of CNVs”). The fixed factors of the model included: diagnoses (AD and control), chromosomes (“Chr”, 22 autosomes as categorical variable), sex (male and female), brain regions (“Brain reg”, temporal cortex, hippocampus CA1, hippocampus CA3, cerebellum, entorhinal cortex) and coverage. Considering correlation due to repeated measurements on the same subject, individual (“Indiv”) effects were added as a random factor. In addition to the interaction of chromosomes and diagnoses (i.e. “Chr * Diagnosis + Coverage”), the effects of the chromosomes were also tested individually (i.e “Chr + Diagnosis + Coverage”. The results were not significant across any of the seven GLMM models tested, in any of the analyses datasets. In the main text, we report the lowest “diagnoses” p-value across the seven tests. Note that we did not apply any multiple testing correction. **E** The heatmap shows the genome-wide copy number profile of cells analyzed in the corrected data (*n* = 10) with at least one reliable CNV. CNVs, brain regions and diagnoses are illustrated in different colors (see the colour key on the left of the figure). Each row shows a cell and each column shows a chromosome.

We also repeated the same analysis on the van den Bos 2016 dataset, from which originally only aneuploidy was reported. Here we identified 11 CNV events across 883 cells in 10 AD individuals and 3 CNV events across 585 cells in 6 controls. The difference was in the same direction as in our dataset, but again not significant (GLMM, p *≥* 0.79) (Fig. 4B, Additional file 2: Table S2, Additional file 3: Table S3).

## Discussion

To the best of our knowledge, this is the first study to use LCM to collect neuronal nuclei for scWGS. Our results showed that LCM-isolated cells suffered from significantly higher within-cell read depth variation compared to FACS-isolated ones. One random source of high variation could be cross-contamination of LCM-isolated cells during the isolation (41), which in turn might be reflected in the down-stream analysis as duplications. In line with this possibility, we found that the number of duplications (*≥*3-somy) is higher than the number of deletions (0- and 1-somy) in the uncorrected data (deletion to duplication ratio: 0.39). We applied several elimination steps to remove “noisy” cells and to filter nominally false positive CNVs. After these elimination steps, the deletion to duplication ratio increased to 6.46 in the uncorrected-filtered data.

In addition to filtering the uncorrected data, we devised a PCA-based denoising approach to remove systematic variation across the genome, which could be experimentally-induced, but could also reflect convergent somatic CNVs also shared among different individuals. Segments systematically deviating from the genome average have also been described in other neuronal scWGS datasets (6). Our results showed that PCA-based denoising can strongly reduce within-cell variance in CN among cells. If the noise that was removed is experimentally-induced, then our result means that this noise was partly shared among cells and not entirely random. One source of systematic bias might be genome-wide variation in the propensity to DNA degradation and/or DNA amplification, perhaps due to GC content, chromatin structure or nuclear location of chromosomal segments (40). Such biases would be shared among cells and effectively removed by PCA.

Beyond the technical biases, biological factors could also explain the higher read-depth variability in LCM-isolated than FACS-isolated neurons. Chronister and colleagues reported that CNV frequencies in neurons (4%–23.1%) are higher than non-neuronal cells (4.7%–8.7%) (6). Moreover, cytological studies suggested that AD brains harbour hyperploid neurons more frequently than healthy controls (10). Consistent with the latter report, we found that neurons from AD patients tend to have higher IOD than control individuals. Also, the cerebellum, which is relatively spared from AD, had lower IOD than the entorhinal cortex and hippocampal areas (but not the temporal cortex). This might be interpreted as a reflection of biological factors on the read-depth variation which is captured efficiently in LCM data. Indeed, if the FACS procedure eliminated cells having abnormal karyotypes, this would result in a cell population with artificially uniform and “clean” ploidy levels. In conclusion, we predict that although random factors (*e*.*g*. contamination) and systematic biases most likely contribute to relatively high variation in LCM-collected scWGS data, biological variation may also be a contributor.

scWGS is a promising method for predicting CNVs with limited sequencing per cell. However, as in our study, within-cell variation that may represent false-positive CNVs hinder analyses in low coverage data. Our PCA-based denoising method can be used as a practical solution for *in silico* cleaning of such data. The approach is based on the idea that somatic CNVs are randomly distributed in the human genome and are particular to each cell. One possible drawback of this approach is that if some neurons from the same individual share the same CNV due to shared developmental ancestry, our method will eliminate such real signals. A more subtle approach could take into account possible clonal relatedness among cells (42). Another drawback could arise if certain genomic regions are predisposed to undergo copy number changes; in that case, our method may cause overcorrection. In our dataset, observing an unexpectedly high frequency of CNVs (23.9 events) per cell in the uncorrected version, we chose to remove *≥* 90% of the common variance. After applying PCA-based correction, the CNV rate per cell decreased by 95.8% (Fig. 3E). This, in turn, resulted in a lower number of CNVs per cell in the corrected data, even compared to published datasets. This difference might be attributable to the overcorrection of normalized read counts.

We note that our PCA-based approach could also be used to detect recurrent breakpoints in single-cell cancer genomics. Because clonal cancer cells would also inherit the same CNVs, shared CNV breakpoints identified in PCAs can be used to study clonal evolution.

Our study has several limitations. First, we only focused on relatively large (*≥*10 Mb) CNVs for sake of sensitivity. However, smaller CNVs may still be much more common and could have contributions to neurodegenerative disease. Future studies on somatic genomic variation in AD might therefore focus on a smaller scale (*<*10 Mb) CNVs, for which improvement of experimental protocols and/or the use of higher coverage data appears to be needed (43). Second, our PCA-based denoising is expected to have removed any CNVs and aneuploidies that are shared among neurons (instead of being cell-specific), due to common origin in the same individual or due to recurrent mutations. Therefore our results only pertain to single cell-specific CNVs. Third, our analysis of published data from van den Bos et al. (2016) could not include a large fraction of cells that they had discarded for showing high depth variability. Finally, recent work has suggested that CNV-bearing neurons may be eliminated through a lifetime in neurotypical individuals (6), and work on hyperploid neurons has also suggested selection against hyperploidy during AD progression (10). This raises the possibility that dynamic elimination may have obscured a possible signal of AD-control difference in neuronal CNV loads, because our sample size did not allow studying disease stage as a separate factor.

## Conclusion

Our main motivation in this study was to describe the relative prevalence of CNVs in the AD brain, where the evidence has been equivocal. Contrary to earlier cytogenetic work, a scWGS study had reported no difference in neuronal aneuploidy levels in the frontal cortex of AD patients versus controls (5). However, the CNV load in different brain regions and relative frequency to the healthy age-matched controls had remained unclear. For example, the entorhinal cortex and hippocampal CA1 have roles in memory formation and learning and are the earliest and most heavily affected regions in AD (44). On the other hand, hippocampal CA3 is less affected, and neurons in the cerebellum are thought to be relatively spared from neurodegenerative disease (34). Here we tackled the same question by comparing AD patients and controls using LCM-isolated cells across five different brain regions, either using the raw data (*n* = 588 cells after filtering) or using a denoising approach (*n* = 1301 cells after filtering). To our knowledge, this is the first dataset that includes scWGS data from pyramidal neurons isolated from AD and control brains in multiple brain regions. Although our AD sample contained slightly higher CNV frequencies than the control sample, none of the comparisons was statistically significant. Our analysis of the van den Bos 2016 dataset yielded a qualitatively similar result, also consistent with the original observation of no significant difference in aneuploidy levels in this dataset (5). Overall, our results call for further research into the possible role of CNVs in AD pathogenesis.

## Methods

### Tissue sources

Frozen postmortem human brain tissues -temporal cortex, hippocampal subfields cornu ammonis (CA) 1, hippocampal subfields cornu ammonis (CA) 3, cerebellum (CB) and entorhinal cortex (EC)- from a total of 13 AD patients and 7 healthy age-matched controls were obtained from the GIE NeuroCEB Brain Bank (France). The Braak stages of the samples were provided by the GIE NeuroCEB Brain Bank. All experiments were conducted at Paul-Flechsig-Institute (Leipzig University, Germany).

### Ethics Statement

The samples were obtained from brains collected in a Brain Donation Program of the Brain Bank NeuroCEB run by a consortium of Patients Associations: ARSEP (association for research on multiple sclerosis), CSC (cerebellar ataxias), LECMA (European league against Alzheimer disease) and France Parkinson. The consents, that have been validated by the Ethical Committee Ile de France 6, were signed by the patients themselves or their next of kin in their name, in accordance with the French Bioethical Laws. The Brain Bank NeuroCEB has been declared at the Ministry of Higher Education and Research and has received approval to distribute samples (agreement AC-2013-1887). The autopsy protocol has been approved by the Biomedicine Agency as requested by the French Law.

### Fluorescence-activated cell sorting (FACS)

Neuronal nuclei were extracted following the protocol described in (45). Briefly, frozen brain samples were thawed in the hypotonic lysis buffer. Neuronal nuclei were stained with propidium iodide and sorted using BD FACSAria II SORP (BD Biosciences). Genomic DNA was then isolated and amplified as described below (see scWGS library preparation and sequencing).

### Laser capture microdissection (LCM)

Frozen brain samples at –80°C were thawed to −20°C, sliced using CryoCut Freezing Microtome at 30 µm thickness, and mounted on a membrane slide (Carl Zeiss). After staining with cresyl violet, single cells were cut out and placed into an adhesive cap by PALM MicroBeam (Carl Zeiss). Neurons of the individual 5603 were collected using both FACS (*n* = 12) and LCM (*n* = 64).

### scWGS library preparation and sequencing

Genomic DNA was amplified using WGA4 (GenomePlex^®^ Single Cell Whole Genome Amplification Kit) and then purified using the MinElute PCR Purification Kit (Qiagen). The specific adapters were added to the DNA via Phusion^®^ PCR followed by purification with the MinElute PCR Purification Kit (Qiagen). Sample quality was evaluated using agarose gel electrophoresis. Sequencing was performed on the HiSeq2500 platform (Illumina) with paired-end 100 bp (PE100) or 150 bp (PE150) modes.

### Read quality control and alignment

The *F astQC* tool (version 0.11.9) was used to check the quality of the raw Illumina reads. The results of *F astQC* were summarized using *MultiQC* (version 1.9) (46). The mean sequence lengths of the reads (ranging between 101 and 151) were inspected using the output of the *MultiQC* (*general*_*stats*_*table*). To avoid biases that would affect the interpretation of the results, all reads were trimmed to a length of 66 (the longest possible length in all reads). Illumina adapter and low-quality bases (the first 35 bp) were removed using *Trimmomatic* (47) with the following parameters: “ILLUMINACLIP:TruSeq3-PE-2.fa:2:30:10:8:TRUE HEADCROP:35 MINLEN:66 CROP:66”. The quality of the trimmed reads was checked again using both *F astQC* and *MultiQC*. Adapter-trimmed paired-end FASTQ files were mapped to the hg19 human reference genome (/ftp://ftp.ensembl.org/pub/release-75/fasta/homo_sapiens/dna/) using **<u>B</u>**urrows-**<u>W</u>**heeler **<u>A</u>**lignment (*BWA* v.0.7.17) (48) with “aln” and “sampe” options.

### Filtering

The output of the *BWA* aligner in **<u>S</u>**equence **<u>A</u>**lignment/**<u>M</u>**ap (SAM) format was further processed by *SAMtools* v1.10 (49) to obtain high-quality uniquely aligned reads. The applied steps are as follows:

1. keep reads mapped in proper pair and discard reads marked with SAM flag 3852:

~~~
samtools view -f 2 -F 3852 -b file.sam > file.bam
~~~
2. extract uniquely mapped reads from BAM files:

~~~
samtools view -h file.bam | egrep -i “^@ | XT:A:U” |
samtools view -Shu - > file.bam2 (39)
~~~
3. obtain reads having MAPQ scores 60:

~~~
samtools view -h -q 60 file.bam2 > file.bam3
~~~
4. sort BAM files:

~~~
samtools sort file.bam3 > file.sorted.bam
~~~
5. filter out PCR duplicates:

~~~
samtools rmdup -S file.sorted.bam file_rm.sorted.bam
~~~
6. index BAM files:

~~~
samtools index -b file_rm.sorted.bam
~~~
7. convert BAM file into BED format using the *Bedtools* “bamToBed” command (*Bedtools* v2.27.1) (50).

### Coverage

*Bedtools* v2.27.1 algorithm “genomeCoverageBed” was used to obtain coverage of the bases on each BAM file. Then the output file was used to calculate the coverage of each sample.

### CNV prediction and cell elimination

CNV calling was performed using *Ginkgo* (35). The command-line version of *Ginkgo* was downloaded from https://github.com/robertaboukhalil/ginkgo. The tool was run under the following settings: (1) variable size of 500 kb bins (39) based on simulations of 76 bp reads aligned with *BWA*, (2) independent segmentation method, (3) ward and euclidean options for the clustering method and clustering distance metric, respectively. Before the segmentation step, GC correction was performed by *Ginkgo* using the *R* function “LOWESS” (see (35)). For segmentation, *Ginkgo* uses the CBS algorithm implemented in DNAcopy in *R* (51). DNAcopy runs with the following parameters: alpha=0.0001, undo.SD=1, min.width=5 (40).

The number of reads was divided into the variable size of 500 kb bins that correspond to 5578 genomic windows. Only cells with *>* 50, 000 reads were kept in downstream analyses (approximately nine reads per window), resulting in *n* = 1337 cells.

### Published datasets

#### The van den Bos 2016 dataset

Data was downloaded from EBI ArrayExpress with the accession numbers E-MTAB-4184 and E-MTAB-4185 (5). Only the cells that were reported as having good quality libraries were included in the analysis (AD:883; control:586; Down’s syndrome:34). Adapter sequences were trimmed with the following parameters: “ILLUMINACLIP:adapter.fa:2:30:10:8:TRUE MINLEN:51”. Single end reads were aligned to the hg19 human reference genome using *BWA* with “aln” and “samse” options. The remaining steps are the same as those described in sections Filtering, except that here we used the SAM flag 3844 (because this dataset was single-end sequenced) and used MAPQ scores 20 (because this dataset did not have enough reads which having the MAPQ 60). Note that due to the missing sample information in the database, the number of analyzed cells in this work does not match what van den Bos and colleagues reported in their original publication.

### The McConnell 2013 dataset

FASTQ files of 110 cells were downloaded from the NCBI SRA database with accession number SRP030642 (36). Adapter sequence was trimmed with the following parameters: “ILLUMINA-CLIP:adapter.fa:2:30:10:8:TRUE MINLEN:39”. Paired-end reads were aligned to the hg19 human reference genome using *BWA* with “aln” and “sampe” options. The remaining steps are the same as those described in sections Filtering.

### Statistical modeling of CNV frequencies and IOD levels

When modelling CNV frequencies, our null hypothesis was no difference in the frequency of CNVs in the AD brain when compared to healthy controls. The overdispersed and zero-dominated nature of the response variable, i.e. the frequency of CNVs, suggested that the data should be fitted using a zero-inflated negative binomial model. For this reason, we used the “glmmadmb” function (package: glmmADMB)(52) in *R* 3.6.3 with the following parameters: “zero-inflated = TRUE” and “family = nbinom1”. The fixed factors of the model were diagnoses (AD and control), chromosomes (autosomes), sex (male and female), brain regions (temporal cortex, hippocampus CA1, hippocampus CA3, cerebellum, entorhinal cortex), and coverage per cell. The individual effect was added as a random factor. Note that sex could not be used as a fixed factor in the van den Bos 2016 dataset because cells that remained after filtering only belonged to females.

When modelling the IOD data, we used the same approach as above. Levels of the response variable, IOD, was predicted using diagnoses (AD and control), brain regions (temporal cortex, hippocampus CA1, hippocampus CA3, cerebellum, entorhinal cortex) and coverage as explanatory variables using the “glmmadmb” function (package: glmmADMB) (52) in *R* 3.6.3. Individual effects were added as a random factor. The distribution of the IOD was right-skewed and the model was run with the “family = gamma” parameter.

To compare the IOD across different brain regions, we used “lme” function (package: nlme) in *R* 3.6.3 with diagnoses as fixed effects and the individual as a random effect.

### Copy number statistics

After reads were mapped into the bins, read counts in each bin were divided by the mean read counts across bins for each cell. This value corresponds to the normalized read counts as calculated by *Ginkgo* (see (35)).

A *Z*_1_-score for each CNV was calculated using the normalized read counts. It was calculated as the cell mean (mean normalized read counts across autosomes) minus the CNV mean (mean read counts between CNV boundaries) divided by the standard deviation (sd) of CNV:

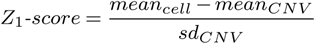

The *Z*_2_-score of each CNV was calculated by calculating the difference between the Ginkgo-estimated integer copy number state (1 or 3) and the observed normalized read count, dividing by the standard deviation (sd) of the normalized read counts:

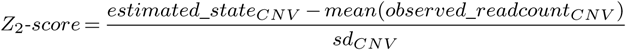

CNVs with two standard deviations below or above the cell’s mean and CNVs with *Z*_2_-score smaller than or equal to 0.5 were kept in the analysis. Using these combinations, monosomy X (≥ 90% of the chromosome’s length) was correctly predicted in 58.1% (217 of 373) of males in the uncorrected data.

### Principal component analysis (PCA)

To remove experimental noise from the data, the following steps were applied for every cell: *(1)* one cell (x) at a time was discarded from the analysis. For the remaining cells, PCA was applied on the normalized read counts using the “prcomp” function with the parameter “scale.=TRUE” in *R* 3.6.3. *(2) n* PCs that explained at least 90% of the variance in total was chosen. *(3)* To remove the effect of the chosen PCs from the focal cell x, a linear regression model with normalized read counts from cell x as a response, and the *n* PCs as explanatory variables was constructed using the *R* “lm” function. *(4)* Residuals from this model were calculated. *(5)* To prevent errors during a lowess fit of GC content (log transformation of negative residuals produces NaNs), plus one was added to the residuals. If there still remained values less than or equal to zero, those values were replaced with the smallest positive number for the focal cell x. *(6)* The resulting value was set as a new value of the focal cell x, and *Ginkgo* was run with the new values.

Also, PCA of the normalized read counts across different datasets was performed in *R* 3.6.3 using the “prcomp” function with the parameter “scale.=FALSE”.

## Author’s contributions

MS, IY, UU, TA, CD conceived and supervised the study. SLT and CD prepared brain tissue and performed neuropathological diagnosis. VR, JB and SKW carried out the LCM and library preparation. ZGT performed analyses with support from PP, EY, and UI. ZGT and MS wrote the manuscript. All authors discussed the results and edited the manuscript.

## Acknowledgements and funding

We thank all colleagues at the METU CompEvo group, in particular, Sinan Can Açan for helpful suggestions, ?Ismail Sag?lam for discussion, and Kıvılcım Başak Vural for their support and help. Funding: The work was supported by an ERA.NetRus grant “significans” (coordinator: Thomas Arendt [FKZ:DLR/01DJ16018], partners: Ivan Iourov, Mehmet Somel, Charles Duyckaerts). The project was funded by a grant from Alzheimer Forschung Initiative (AFI; 17021) to U.U. and TÜB?ITAK grant 215Z495. ZGT was supported by a YOK PhD fellowship. Figure 1A and Figure 4A were created with https://biorender.com/.

## Availability of supporting data

All data from this study have been submitted to the European Nucleotide Archive (ENA) repository under accession number PRJEB51941. The codes and additional information can be found in the Github repository https://github.com/zgturan/brain_CN.

## Additional Files

**Additional file 1 — Table S1**. Sample information.

**Additional file 2 — Table S2**. Information about the cells and significant CNVs. **A-B**: Uncorrected data. **C-D** corrected data. **E-F**: van den Bos

**Additional file 3 — Table S3**. Tables show the GLMM results across different datasets and models. Note that we did not apply any multiple testing correction. **A-G**: Uncorrected filtered data. **H-N**: PCA-corrected data. **O-Q**: van den Bos. **R-X**: Uncorrected data.

**Additional file 4. Figures S1**: Information about the samples of uncorrected and PCA-corrected data. **Figures S2**: CN profiles of three cells deviating from the range of 1.9 and 2 after correction. **Figures S3**: Examples of CNVs that failed to pass and passed the filtering criteria in the uncorrected-filtered data and PCA-corrected data.

