## Additional file 4 for "Somatic copy number variant load in neurons of healthy controls and Alzheimer’s disease patients"

### Supplemental Information

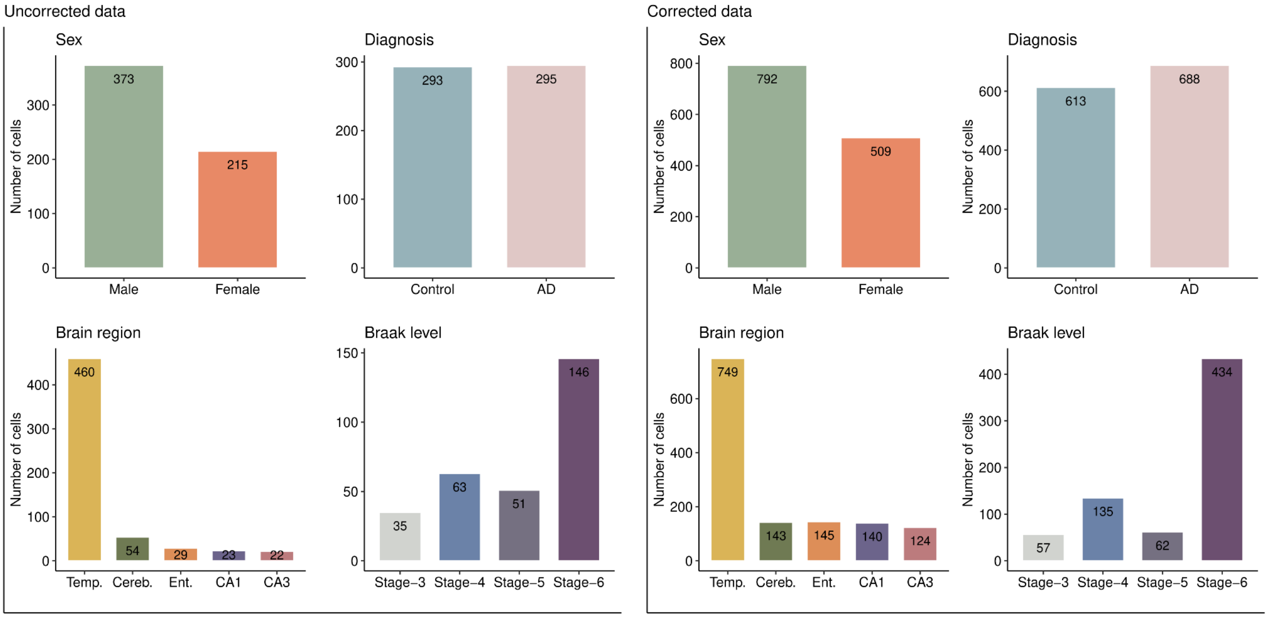


**Figure S1.** Information about the samples of uncorrected and PCA-corrected data. Distribution of samples according to sex (female, male) (*A*), diagnoses (Alzheimer's Disease, control) (*B*), brain regions (temporal cortex, cerebellum, entorhinal cortex, hippocampal subfields CA1, hippocampal subfields CA3) (*C*) and Braak level (stage 3, 4, 5 and 6) (*D*).

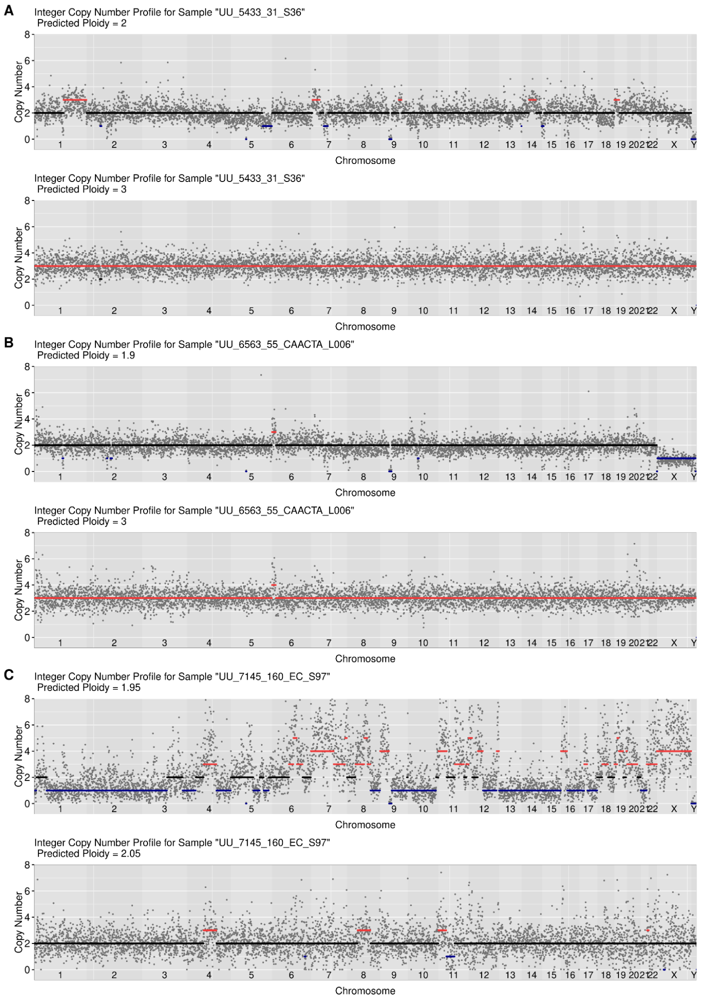


**Figure S2.** CN profiles of three cells deviating from the range of 1.9 and 2 after correction. The upper panels show the uncorrected data, and the lower panels show the corrected data. X-axes show chromosomes and Y-axes illustrate the CN profile of chromosomes estimated by *Ginkgo*. Each grey dot represents the scaled read counts per bin. Amplifications (CN>2) are shown in red; deletions (CN<2) in blue; disomy (CN=2) in black.

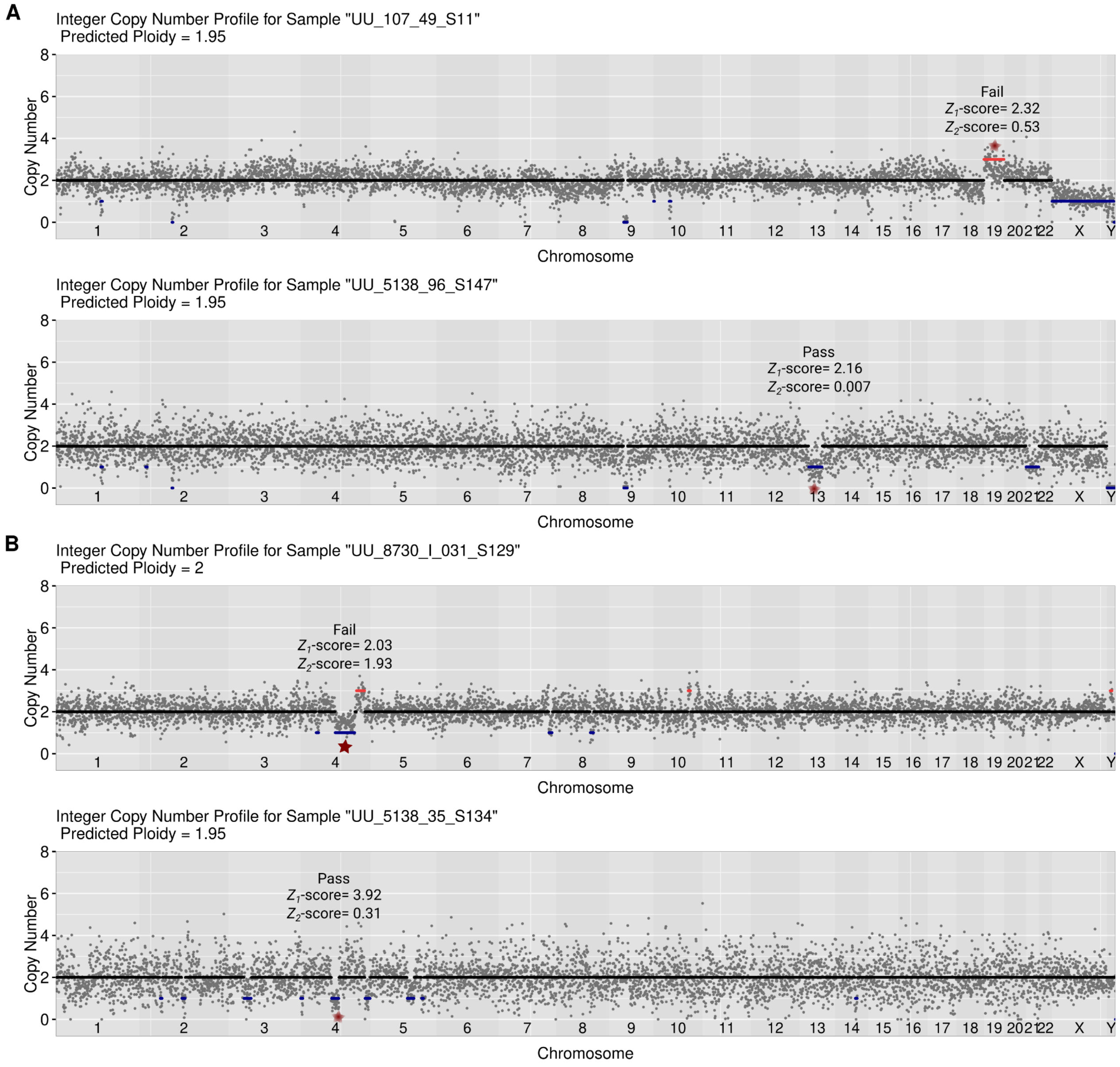


**Figure S3.** Examples of CNVs that (upper panels) failed to pass and (lower panels) passed the filtering criteria ($Z_1$-scores ≥2 and $Z_2$-scores ≤0.5) in the uncorrected-filtered data (*A*) and PCA-corrected data (*B*). Absolute values of the $Z_1$- and $Z_2$ scores for the CNV are indicated on the plot. The CNVs in question (that failed or passed the filtering) are marked with red stars. X-axes show chromosomes and Y-axes illustrate the CN profile of chromosomes estimated by *Ginkgo*. Each grey dot represents the scaled read counts per bin. Amplifications (CN>2) are shown in red; deletions (CN<2) in blue; disomy (CN=2) in black.
